# Using Y chromosomal haplogroups in genetic association studies and suggested implications

**DOI:** 10.1101/048504

**Authors:** A. Mesut Erzurumluoglu, Denis Baird, Tom G. Richardson, Nicholas J. Timpson, Santiago Rodriguez

## Abstract

Y chromosomal (Y-DNA) haplogroups are more widely used in population genetics than in genetic epidemiology, although associations between Y-DNA haplogroups and several traits (including cardio-metabolic traits) have been reported. In apparently homogeneous populations, there is still Y-DNA haplogroup variation which will result from population history. Therefore, hidden stratification and/or differential phenotypic effects by Y-DNA haplogroups could exist. To test this, we hypothesised that stratifying individuals according to their Y-DNA haplogroups before testing associations between autosomal SNPs and phenotypes will yield difference in association. For proof of concept, we derived Y-DNA haplogroups from 6,537 males from two epidemiological cohorts, ALSPAC (N=5,080, 816 Y-DNA SNPs) and 1958 Birth Cohort (N=1,457, 1,849 Y-DNA SNPs). For illustration, we studied well-known associations between 32 SNPs and body mass index (BMI), including associations involving *FTO* SNPs. Overall, no association was replicated in both cohorts when Y-DNA haplogroups were considered and this suggests that, for BMI at least, there is little evidence of differences in phenotype or gene association by Y-DNA structure. Further studies using other traits, Phenome-wide association studies (PheWAS), haplogroups and/or autosomal SNPs are required to test the generalisability of this approach.

## Introduction

The interpretation of genetic association studies (including candidate gene studies and genome-wide association studies, GWAS) requires consideration of issues including population stratification, gene-gene interaction and gene-environment interaction [1-3]. The relevance of these factors and in particular population structure and haplotype background [4], has been explored by the analysis of autosomal markers. In contrast, non-recombining genetic variation such as Y chromosomal (Y-DNA) haplogroups, has rarely been considered in the design and interpretation of genetic association studies-although there are examples including direct testing of the association between Y-DNA haplogroups and phenotypes, including cardio-metabolic diseases [5-10].

Analyses of the non-recombining regions of the Y chromosome (NRY) in different populations provide genealogical and historical information [11, 12]. Y chromosomal lineages, through the analysis of short tandem repeats (STR), have proven useful when determining whether two apparently unrelated individuals descend from a common ancestor in recent history (<20 generations). However, using of modern genotyping arrays coupled with extensive and publicly available SNP data, researchers now possess the ability to identify which ancient ethnic group to one’s paternal ancestor belonged to. Comprehensive single nucleotide polymorphism (SNP) data also enabled the publication of well-established Y-DNA haplotypes and constantly updated phylogenetic trees [13-15]. This is why genetic variation in this uniparentally inherited chromosome can be used to define groups of Y-DNA haplotypes which share a common ancestor with a SNP mutation.

Haplogroups derived from Y-chromosomal variation can be used to provide information about the paternal ancestry of an individual and population genetic events (e.g. migrations, bottle necks) [16-18]. Phylogenetic relationships between haplogroups are well known and there is wide-spread knowledge of the frequency and the type of haplogroups present in almost all geographical regions throughout the world. For example the Y-DNA haplogroup R1b1 is frequent in Europe and infrequent or absent in other continents/sub-continents.

Facilitating the process of linking haplotype assignment to GWAS studies there is comprehensive information about the SNPs which define each haplogroup, approximate time and (most probable) region of origin, (current) area of highest frequency and the most prevalent (ancient) haplogroup present in different regions.

Even for homogeneous populations (according to autosomal SNPs), there is underlying Y-DNA haplogroup variation. We have previously analysed Y-DNA haplotypes in a large epidemiological cohort in relation to confounding by genetic subdivision [19].

In the present work, we stratify groups of individuals according to their Y-DNA haplogroups to (i) test for presence of additional structure due to Y-DNA haplogroup variation having taken into account principal component analysis (PCA) using autosomal markers and if there is (ii) test if this additional structure has any potential confounding effects on genetic association studies (e.g. direct association, epigenetic, epistasis). As a proof of concept, we chose to study the association between 32 common SNPs which are known to be reliably associated with BMI. The use of these common genetic variants limits our analyses to the largest BMI effect loci.

## Methods

### Participants and Ethics

The Avon Longitudinal Study of Parents and Children (ALSPAC) is a longitudinal, population-based birth cohort study that initially recruited >13,000 pregnant women residing in Avon, United Kingdom, with expected dates of delivery between April 1, 1991 and December 31, 1992. There were 14,062 liveborn children. The study protocol has been described previously [20, 21] and further details are available on the ALSPAC website (http://www.bris.ac.uk/alspac). Please note that the study website contains details of all the data that is available through a fully searchable data dictionary (http://www.bris.ac.uk/alspac/researchers/data-access/data-dictionary/).

Height and weight measurements were performed on children who attended a 9 years focus group clinic [mean age of participant 9 (± 0.32 years)]. Ethical approval for all aspects of data collection was obtained from the ALSPAC Law and Ethics Committee (institutional review board 00003312). Written informed consent for the study was obtained for genetic analysis.

The National Child Development Study (NCDS), otherwise known as the 1958 British birth cohort (1958BC), started as a perinatal mortality and morbidity survey looking at all births in England, Wales and Scotland in a single week in 1958. This included an original sample of 17,638 births (in addition to a further 920 immigrants born in the same reference week). Cohort members were further followed-up by medical examinations (at 7, 11 and 16 years of age) and interviews (at ages 23, 33 and 42). The first biomedical assessment was conducted between September 2002 and March 2004 by trained nurses from the National Centre for Social Research, who visited the homes of cohort members at age 44-45 years [22].

### Genotyping and Imputation

#### ALSPAC

A total of 9,912 participants were genotyped using the Illumina HumanHap550 quad genome-wide SNP genotyping platform by Sample Logistics and Genotyping Facilities at the Wellcome Trust Sanger Institute and LabCorp (Laboratory Corporation of America). PLINK software (v1.07) was used to carry out quality control (QC) measures [23]. Individuals were excluded from further analysis on the basis of having incorrect sex assignments, minimal or excessive heterozygosity (< 0.320 and > 0.345 for the Sanger data and < 0.310 and > 0.330 for the LabCorp data), disproportionate levels of individual missingness (> 3%), evidence of cryptic relatedness (> 10% IBD) and being of non-European ancestry (as detected by a multidimensional scaling analysis seeded with HapMap 2 individuals). Autosomal SNPs with a minor allele frequency of < 1% and call rate of < 95% were removed. Furthermore, only autosomal SNPs which passed an exact test of Hardy-Weinberg equilibrium (P > 5x10^−7^) were considered for analysis. After QC, 8,365 unrelated individuals who were genotyped at 500,527 autosomal SNPs were available for analysis. Known autosomal variants were imputed with MACH 1.0.16 Markov Chain Haplotyping software [24, 25], using CEPH individuals from phase 2 of the HapMap project (hg18) as a reference set (release 22) [26].

#### 1958 Birth Cohort (1958BC)

3000 individuals were genotyped on the Illumina 1.2M chips [Dataset ID: EGAD00000000022]. QC measures were as described above. No imputation was carried out as rs8050136) was the only SNP analysed in the 1958BC.

### Y-DNA haplogroup determination

For Y-DNA haplogroup determination in ALSPAC, the Y-chromosomal SNPs of all 5,085 male participants in the dataset were used. The pseudo-autosomal SNPs were removed using the PLINK software package [23]. The resulting Y chromosomal genotype (816 SNPs) of each individual was then piped in to the YFitter (v0.2) software (maps genotype data to the Y-DNA phylogenetic tree built by Karafet *et al* [14], available online at sourceforge.net/projects/yfitter) and their respective Y-DNA haplogroup was determined. After removal of individuals with ‘False’ haplogroup determinations (i.e. ones which did not have enough SNPs to reliably determine haplogroup), we were left with 5,080 individuals. Remaining individuals with a haplogroup result which began with the letter R (e.g. R1b1) were clustered in to a single group named ‘Clade R’ and likewise the same was done with the haplogroups beginning with the other letters. The same procedure was carried out in 1958BC and 1,453 male participants’ haplogroups were determined. Only the major haplogroups R and I were considered in the analyses, since there was not enough power for the less frequent haplogroups.

### Association study between Y-DNA haplogroups and BMI

To check for association between BMI and the Y-DNA haplogroups in ALSPAC, a linear regression analysis was carried out using haplogroup R as a baseline (coded 0) and coding haplogroup I as 1. Age and age^2^ were used as covariates in the model. The analysis was repeated in the 1958BC. Production of summary statistics for the two cohorts and all regression analyses were carried out in the STATA statistical package.

### Analysis of the effects of Y-DNA haplogroup on SNPs associated with BMI

In order to study whether well-established associations are still present and/or observable within each Y-DNA haplogroup and whether the effect sizes of the SNPs stayed stable across haplogroups, we analysed 32 SNPs previously reported to be associated with BMI [27]. This enables the analysis of common genetic variation involved in the largest effect sized observed for BMI. All individuals with missing and/or incorrectly measured data were excluded. Individuals with ‘False’ haplogroups (as determined by YFitter) were also removed. At the end of the QC procedure, 2,800 individuals had complete haplogroup, BMI and genotype data. Finally individuals belonging to haplogroups with frequencies less than 1% were also excluded.

BMI data did not follow a normal distribution and inverse rank transformation was used to correct this. SNP dosage values were determined using the software package MaCH [24, 25]. A linear regression analysis between BMI and each of the 32 SNPs was carried out using STATA controlling for age, age^2^ and the first 10 PCs determined by the EIGENSTRAT software [28]. We looked at the PCA adjusted data only in order to see the Y-chromosome sub-structure. We checked for normal distribution of BMI within the two most frequent Y-DNA haplogroup clades observed (i.e. R and I); and also confirmed that the allele frequencies of the autosomal SNPs analysed were similar across the haplogroups. A subgroup analysis was carried out within the Y-DNA haplogroups R and I. any possible interaction between genotype and Y-DNA haplogroup in the analyses were assessed using a likelihood ratio test to compare the two regression models, one which was adjusted for the covariates abovementioned and the Y-DNA haplogroup and another which additionally included an interaction term (i.e. genotype x Y-DNA haplogroup). Although low powered compared to the interaction test mentioned above, a heterogeneity test (i.e. z-test) was carried out across the two haplogroups to check whether there was a difference in effect size (beta coefficient) of all 32 SNPs tested.

## Results

Figure 1a presents the Y-DNA haplogroups observed in ALSPAC. Haplogroup R is the most frequent (72%) and I the second most common Y-DNA haplogroup (19%). Y-DNA haplogroups subclades observed in ALSPAC are shown in Figure 1b. Most of the males in ALSPAC belong to the R1b1b2 haplogroup (over 3,400 individuals) which is also one of the most common haplogroups in Europe (see www.eupedia.com/europe/origins_haplogroups_europe.shtml). Figures 2a and 2b present the Y-DNA haplogroup profile of 1958BC. Five Y-DNA haplogroup were observed. Similar to ALSPAC, haplogroup R was the most frequent (74%), followed by haplogroup I (20%).

**Figure 1a:**
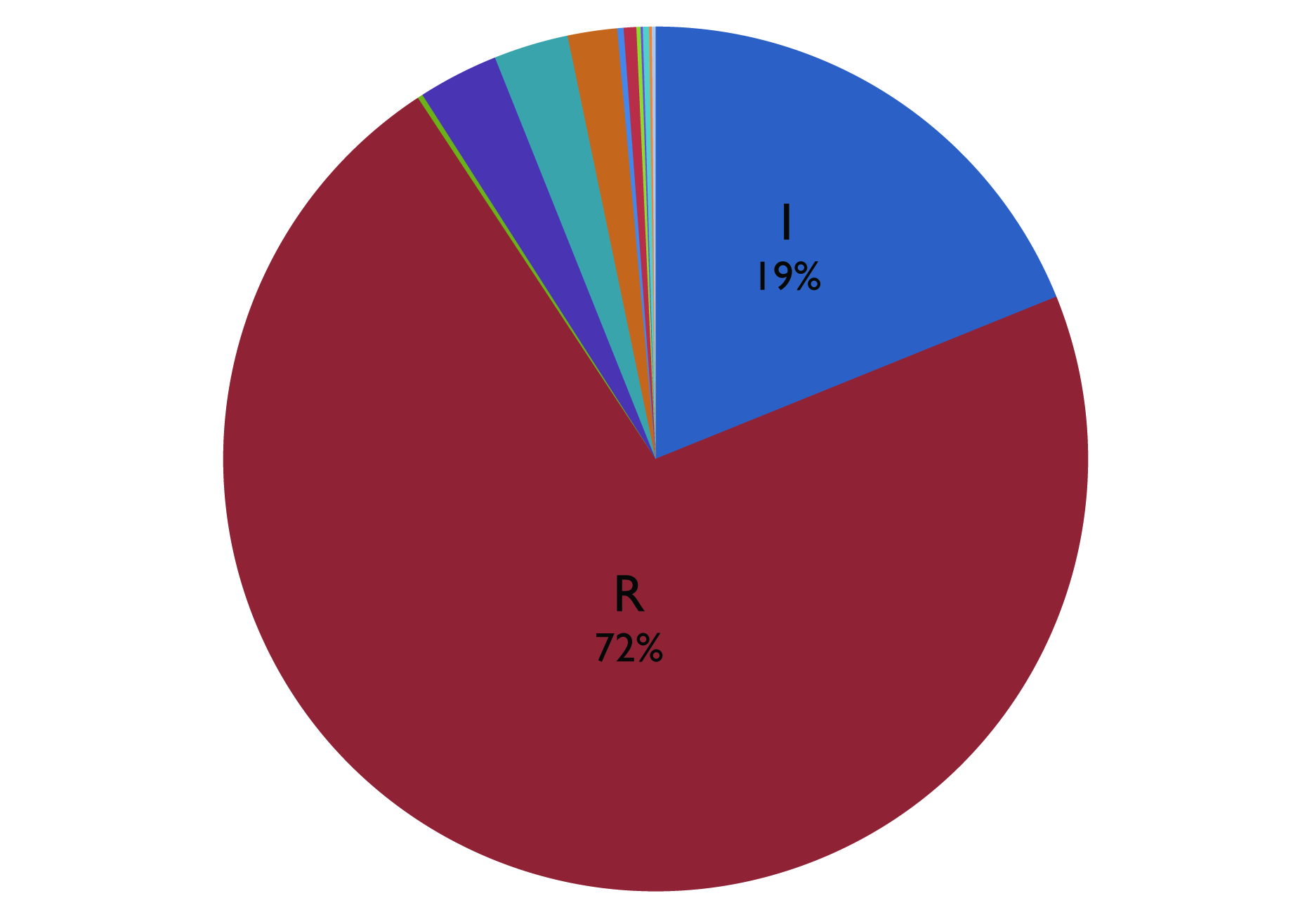
Y-DNA haplogroups in ALSPAC. The ALSPAC cohort has within it individuals belonging to 12 of the major Y-DNA haplogroups (C, E, G, H, I, J, L, N, O, Q, R, T), albeit only 5 of the groups have 50 (>1%) or more individuals in them. These five clades are E, G, I, J and R and have 153 (3%), 94 (1.9%), 960 (19%), 142 (2.8%) and 3564 (72%) individuals in them.

**Figure 1b:**
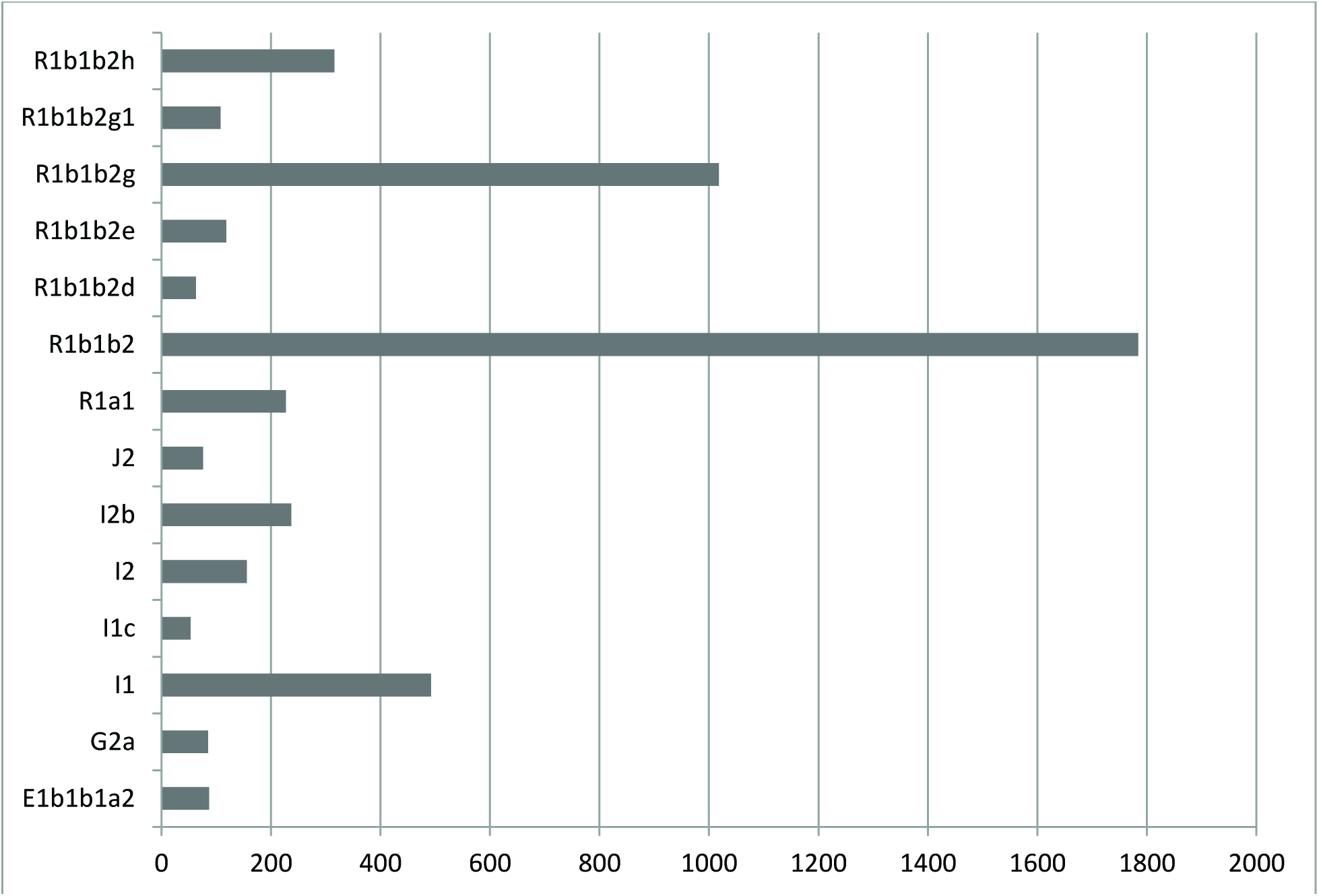
Y-DNA haplogroup frequencies in ALSPAC. Many of the individuals had extensive Y-DNA SNP data (which passed QC), which enabled us to pinpoint exactly which haplogroup they belonged to. Fig. 2a shows the most detailed haplogroup determination; and only the ones with over 50 individuals (>1%) are shown. Where the haplogroup branching halts is an indication of how far we could reliably determine the Y-DNA phylogenetic branch an individual belongs to.

**Figure 2ab:**
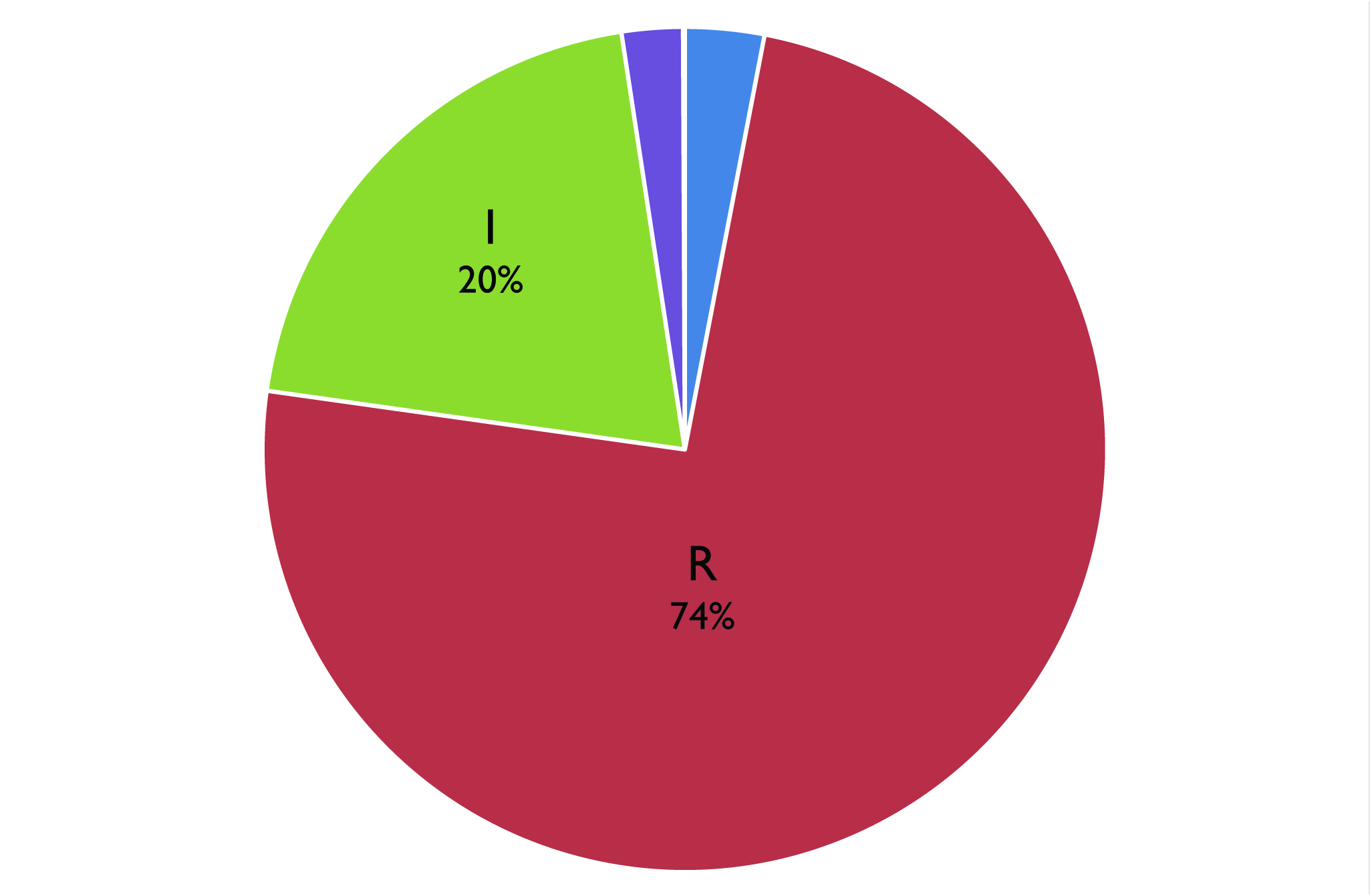
Y-DNA haplogroups in 1958 Birth Cohort (Dataset: EGAD00000000022) The 1958 Birth cohort has within it individuals belonging to five major Y-DNA haplogroups (E, I, J, C, R), albeit 2 of the groups have less than 50 individuals in them. The clades E, I, J, C and R have 44 (3%), 296 (20%), 34 (2%), 1 and 1078 (74%) individuals in them.

**Figure 2b:**
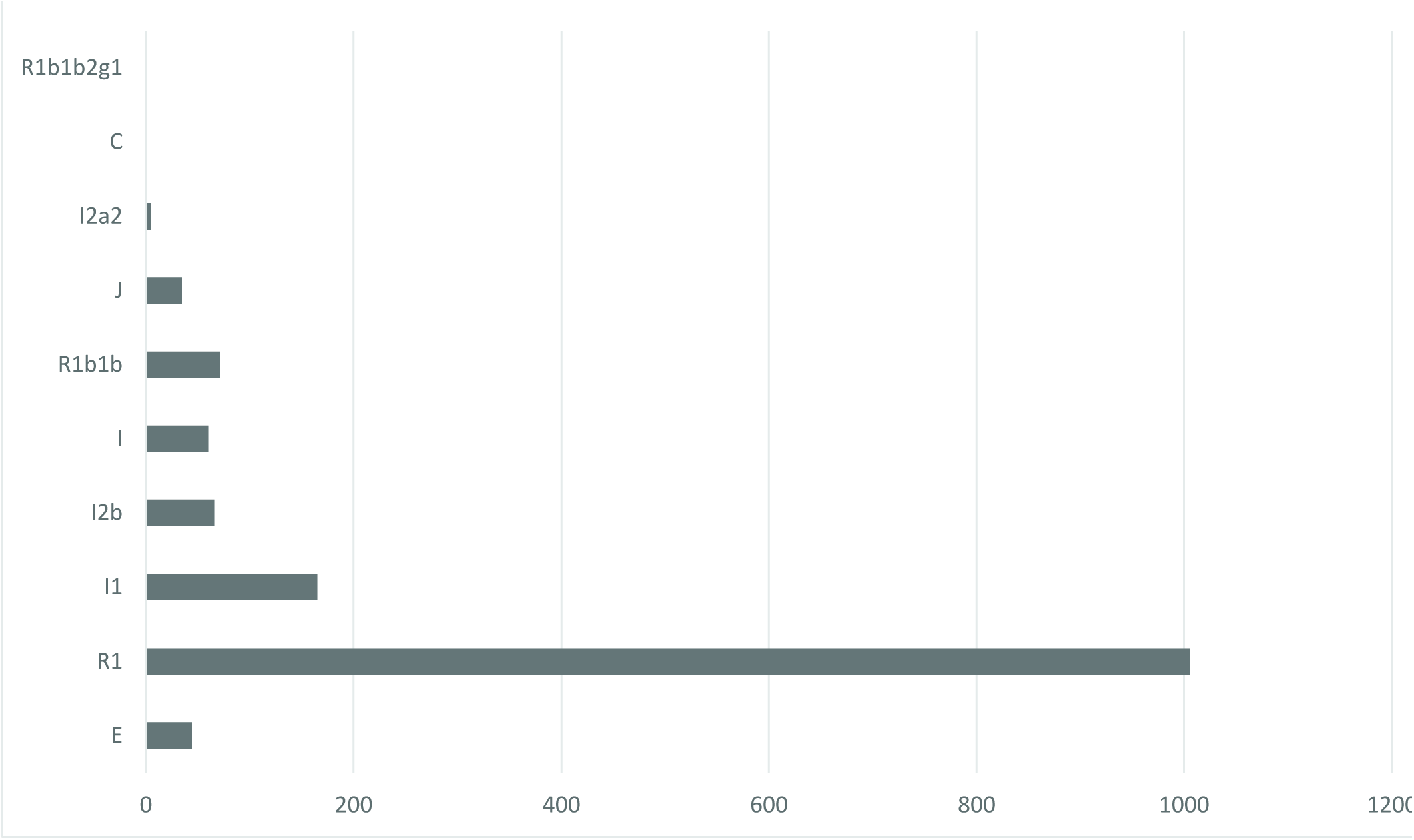
Y-DNA haplogroup frequencies in 1958BC (Dataset: EGAD00000000022) Similar to Fig. 2a, 1958BC provides dense SNP data which enabled deeper haplogroup determination. The frequencies of haplogroups are 1, 1, 5, 34, 71, 60, 66, 165, 1006 and 44 from top to bottom.

There was no strong evidence of association between Y-DNA haplogroups and BMI (P=0.066) in ALSPAC (Table 1) and in the 1958 cohort (P=0.107) (Table 1). Summary statistics of the BMI observed for the two main Y-DNA haplogroups in ALSPAC and 1958BC can be found in Table 2.

**Table 1:**
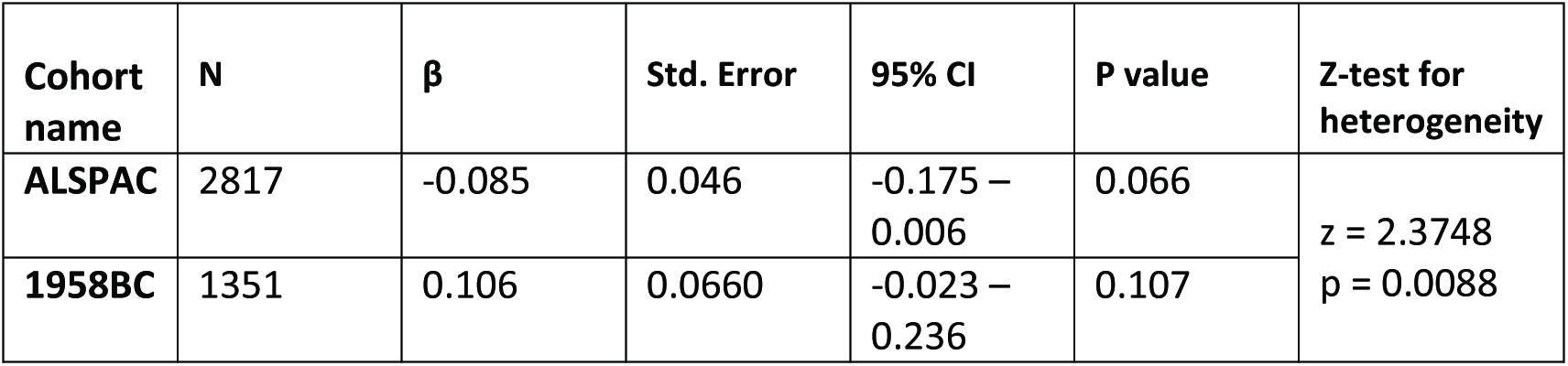
Linear regression between BMI and Y-DNA haplogroup I in two cohorts - ALSPAC and 1958BC. The z-test for heterogeneity shows that the effect size of Y-DNA haplogroup I on BMI is differential depending on the cohort.

**Table 2:** Summary statistics of the two Y-DNA haplogroups for BMI in ALSPAC

| Variable | N | Mean age<br>(range) | Mean<br>BMI | Std.<br>Dev | Min BMI | Max BMI | Std. Error | 95% CI for Mean |
| --- | --- | --- | --- | --- | --- | --- | --- | --- |
| Y-DNA I –<br>ALSPAC | 583 | 7.57<br>(7.07-9.49) | 16.02 | 2.018 | 12.36 | 28.28 | 0.084 | 15.86 - 16.19 |
| Y-DNA R -<br>ALSPAC | 2234 |  | 16.12 | 1.843 | 11.78 | 27.28 | 0.039 | 16.04 - 16.20 |
| Y-DNA I –<br>1958BC | 293 | 23.0<br>(22.9-23.1) | 23.32 | 2.981 | 18.02 | 39.20 | 0.174 | 22.98 - 23.66 |
| Y-DNA R –<br>1958BC | 1058 |  | 23.00 | 2.790 | 14.00 | 37.32 | 0.086 | 22.83 - 23.17 |

Table 3a includes 32 SNPs previously reported to be associated with BMI [27] and presents the association between each SNP and BMI observed for individuals belonging to Y-DNA haplogroups I and R in ALSPAC. Table 3a also shows the results from the heterogeneity test (i.e. z-test) used to compare the effect sizes derived from the two Y-DNA haplogroups. Only one instance of heterogeneity between the two haplogroups was observed (*FTO*) after adjusting for a Bonferroni correction. The highest difference was observed for SNP rs8050136 in *FTO* which yielded a heterogeneity test p value of 0.005 (z heterogeneity test). The likelihood ratio test for interaction between the *FTO* SNP rs8050136 and haplogroup I yielded a p value of 0.008 (Figure 3a). In ALSPAC, there was a difference in the effect size of this SNP within haplogroup I (P=7.00 x 10^−5^, beta=0.266, SE=0.066, N=508) compared with haplogroup R (P=1.4 x 10^−2^, beta=0.079, SE=0.032, N=1,965).

**Figure 3a-c:**
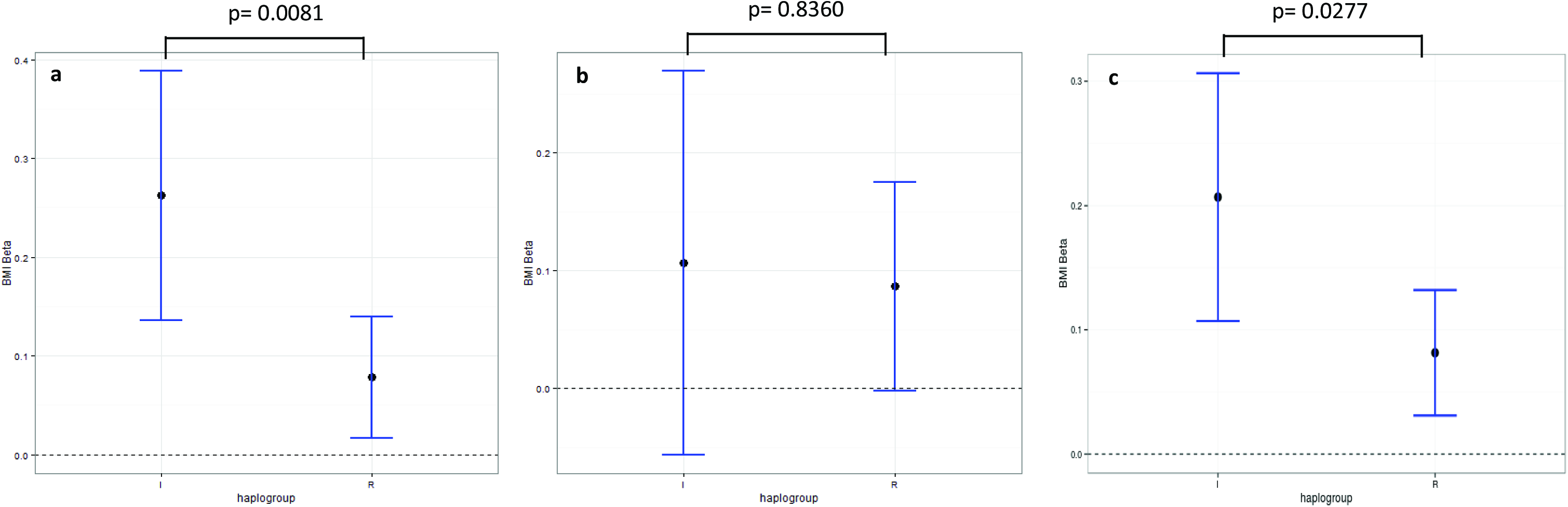
Subgroup analysis comparing effect size of rs8050136 on BMI in two Y-DNA haplogroups. Results from a) ALSPAC b) 1958BC c) ALSPAC and 1958BC combined. The statistics above represent p values from the likelihood ratio test for interaction between Y-DNA haplogroup I and rs8050136. Heterogeneity tests (z test) comparing Y-DNA haplogroups I and R yielded p values of 0.005, 0.4169 and 0.014 for a, b and c respectively. ggplot2 package in R was used to create the plot.

**Table 3a:** Comparison of associations observed between ALSPAC individuals with Y haplogroup I and Y haplogroup R for SNPs previously reported to be associated with BMI (only one of the *FTO* SNPs is shown, see Supp. Table S2 for details).

|  |  | Haplogroup I<br>(508 Individuals) |  |  | Haplogroup R<br>(1965 Individuals) |  |  | Test of Heterogeneity<br>(Z test p value) |
| --- | --- | --- | --- | --- | --- | --- | --- | --- |
| SNP ID | Nearby Gene | P-Value | B-Coef. | St. Error | P-Value | B-Coef. | St. Error |  |
| rs8050136 | *FTO* | 5.45E-05 | **0.261** | 0.064 | 1.60E-02 | **0.076** | 0.032 | **p=0.0050** |
| rs2815752 | *NEGR1* | 7.48E-01 | 0.021 | 0.064 | 4.54E-01 | 0.023 | 0.031 | p=0.9776 |
| rs1514175 | *TNNI3K* | 4.36E-01 | 0.050 | 0.064 | 4.25E-01 | 0.025 | 0.031 | p=0.7252 |
| rs1555543 | *PTBP2* | 3.92E-01 | 0.058 | 0.068 | 2.25E-01 | 0.039 | 0.032 | p=0.8004 |
| rs543874 | *SEC16B* | 3.00E-03 | -0.230 | 0.077 | 1.10E-02 | -0.097 | 0.038 | p=0.1214 |
| rs2867125 | *TMEM18* | 1.50E-02 | 0.207 | 0.085 | 1.00E-03 | 0.130 | 0.040 | p=0.4124 |
| rs713586 | *RBJ* | 4.10E-01 | 0.054 | 0.066 | 4.60E-02 | 0.061 | 0.031 | p=0.9235 |
| rs887912 | *FANCL* | 6.18E-01 | -0.036 | 0.071 | 1.69E-01 | 0.046 | 0.033 | p=0.2949 |
| rs2890652 | *LRP1B* | 1.20E-01 | 0.138 | 0.089 | 2.42E-01 | 0.047 | 0.041 | p=0.3531 |
| rs13078807 | *CADM2* | 1.58E-01 | 0.108 | 0.077 | 4.04E-01 | -0.032 | 0.039 | p=0.1048 |
| rs9816226 | *ETV5* | 1.65E-01 | 0.123 | 0.088 | 6.00E-02 | -0.077 | 0.041 | p=0.0394 |
| rs10938397 | *GNPDA2* | 5.51E-01 | -0.039 | 0.065 | 3.00E-03 | -0.092 | 0.031 | p=0.4617 |
| rs13107325 | *SLC39A8* | 8.83E-01 | 0.018 | 0.122 | 7.00E-02 | -0.106 | 0.058 | p=0.3587 |
| rs2112347 | *FLJ35779* | 3.59E-01 | -0.064 | 0.069 | 1.10E-01 | -0.052 | 0.033 | p=0.8753 |
| rs4836133 | *ZNF608* | 4.97E-01 | -0.047 | 0.068 | 9.25E-01 | -0.003 | 0.031 | p=0.556 |
| rs206936 | *NUDT3* | 4.01E-01 | -0.070 | 0.083 | 9.86E-01 | -0.001 | 0.039 | p=0.4518 |
| rs987237 | *TFAP2B* | 5.23E-01 | -0.056 | 0.087 | 3.80E-02 | -0.082 | 0.039 | p=0.7851 |
| rs10968576 | *LRRN6C* | 3.90E-02 | -0.143 | 0.069 | 7.82E-01 | 0.009 | 0.033 | p=0.0469 |
| rs4929949 | *RPL27A* | 5.46E-01 | 0.040 | 0.066 | 9.68E-01 | 0.001 | 0.031 | p=0.5928 |
| rs10767664 | *BDNF* | 9.72E-01 | 0.003 | 0.078 | 1.36E-01 | 0.055 | 0.037 | p=0.547 |
| rs3817334 | *MTCH2* | 5.63E-01 | -0.040 | 0.069 | 8.63E-01 | 0.005 | 0.031 | p=0.5519 |
| rs7138803 | *FAIM2* | 8.74E-01 | 0.010 | 0.065 | 9.00E-03 | 0.085 | 0.033 | p=0.3036 |
| rs4771122 | *MTIF3* | 5.83E-01 | -0.044 | 0.079 | 4.90E-02 | -0.074 | 0.038 | p=0.7322 |
| rs11847697 | *PRKD1* | 5.49E-01 | -0.106 | 0.176 | 1.00E-03 | -0.234 | 0.072 | p=0.5009 |
| rs10150332 | *NRXN3* | 7.64E-01 | 0.023 | 0.076 | 8.36E-01 | -0.008 | 0.038 | p=0.7152 |
| rs2241423 | *MAP2K5* | 8.50E-02 | -0.130 | 0.075 | 8.18E-01 | 0.009 | 0.038 | p=0.0983 |
| rs12444979 | *GPRC5B* | 9.67E-01 | 0.004 | 0.094 | 2.00E-03 | 0.133 | 0.043 | p=0.212 |
| rs7359397 | *SH2B1* | 5.20E-02 | -0.127 | 0.065 | 6.02E-01 | -0.016 | 0.031 | p=0.1232 |
| rs571312 | *MC4R* | 1.99E-01 | 0.099 | 0.077 | 2.00E-03 | 0.116 | 0.037 | p=0.8423 |
| rs29941 | *KCTD15* | 2.33E-01 | 0.087 | 0.073 | 1.14E-01 | -0.051 | 0.032 | p=0.0834 |
| rs2287019 | *QPCTL* | 4.79E-01 | 0.056 | 0.078 | 8.01E-01 | -0.010 | 0.040 | p=0.4515 |
| rs3810291 | *TMEM160* | 2.18E-01 | 0.099 | 0.080 | 2.57E-01 | 0.042 | 0.037 | p=0.5178 |

**Table 3b:** Comparison of the associations observed between 1958BC individuals with Y haplogroup I and Y haplogroup R for the only SNP showing a P~0.05 in any of the two cohorts analysed.

|  |  | Haplogroup I<br>(293 Individuals) |  |  | Haplogroup R<br>(1058 Individuals) |  |  | Test of<br>Heterogeneity |
| --- | --- | --- | --- | --- | --- | --- | --- | --- |
| SNP ID | Nearby<br>Gene | P-Value | B-Coef. | St. Error | P-Value | B-Coef. | St. Error |  |
| rs8050136 | *FTO* | 0.201 | 0.106 | 0.083 | 0.056 | 0.086 | 0.045 | p= 0.4169 |

The p value for heterogeneity, (as measured by the likelihood ratio test) between Y-DNA haplogroups I and R in relation to the association between *FTO* and BMI, was p= 0.008. The observed results were consistent for all nine *FTO* SNPs, with heterogeneity between Y-DNA haplogroups I and R in all cases (Table 4).

**Table 4:** Comparison of associations observed between ALSPAC individuals with Y haplogroup I and Y haplogroup R for *FTO* SNPs observed in ALSPAC. All z-test p values are less than 0.01.

| SNP ID | Haplogroup I<br>(521 Individuals) |  |  | Haplogroup R<br>(2011 Individuals) |  |  | Test of Heterogeneity |
| --- | --- | --- | --- | --- | --- | --- | --- |
|  | P-Value | B-Coef. | St. Error | P-Value | B-Coef. | St. Error |  |
| **rs8050136** | 5.45E-05 | 0.261 | 0.064 | 1.60E-02 | 0.076 | 0.032 | I<sup>2</sup>=84.9% |
| **rs9940128** | 1.11E-04 | 0.252 | 0.065 | 2.30E-02 | 0.072 | 0.032 | I<sup>2</sup>=83.8% |
| **rs9939609** | 1.11E-04 | 0.252 | 0.065 | 2.30E-02 | 0.072 | 0.032 | I<sup>2</sup>=83.8% |
| **rs9930506** | 8.49E-05 | 0.266 | 0.067 | 2.20E-02 | 0.075 | 0.033 | I<sup>2</sup>=84.7% |
| **rs17817449** | 1.12E-04 | 0.251 | 0.065 | 3.10E-02 | 0.068 | 0.032 | I<sup>2</sup>=84.3% |
| **rs7193144** | 1.11E-04 | 0.252 | 0.065 | 2.00E-02 | 0.074 | 0.032 | I<sup>2</sup>=83.4% |

To test whether the possible differential effect of the *FTO* SNPs replicate in another cohort, the top-hit rs8050136 SNP was analysed in the 1958BC and the results are presented in Table 3b. The likelihood ratio test for interaction (and z heterogeneity test) yielded a p> 0.05 (p= 0.836, z-test p= 0.4169, see Figure 3b).

## Discussion

Population stratification is a potential confounder in genetic association studies. Haplotypic variation and sub-clustering can still be present even after accounting for principal components (see reference [4] for an example). Therefore an apparently homogeneous population (defined by principal component analysis) can harbour different subgroups of individuals. In this work, we analysed whether this was the case for Y-DNA haplogroups.

We used the ALSPAC cohort - formed of a relatively homogeneous group of participants - for proof of concept that Y-DNA haplogroup variation is present even after accounting for principal components. We then looked to see whether this variation could confound the genetic association studies related to BMI. In this work, we also present the Y-DNA haplogroup profiles of two cohorts for genetic epidemiological studies - ALSPAC and the 1958BC. Within a homogenous looking population there were individuals belonging to different paternal lineages. We undertook a stratified analysis of Y-DNA haplogroups in ALSPAC. This can be the case especially if the trait is associated to the haplogroup(s). In this study we observed no strong evidence for differences in SNP/BMI association according to Y-DNA haplogroups in either ALSPAC or the 1958BC.

A key aspect about the relevance of the Y-DNA structure is that there can be an effect on the phenotype associations if the structure is also correlated with BMI and if the actual haplogroup interacts directly with the assessed gene variants. Alternatively, other loci would be enough to obscure inference. Our study showed no clear evidence of this correlation and interaction. However, one could argue that the lack of replication could be explained by heterogeneity of both studies (ALSPAC and 1958 cohort), since the former includes children and the latter, adults. Therefore, the differences in betas and interaction effects could be function of differences between cohorts.

A sub-clustering due to Y-DNA haplogroups can be revealed by plotting the Y-DNA haplogroup information versus the top two PCs on a scatter plot (see ALSPAC example on Figure 4). For the ALSPAC cohort, sub-clustering due to Y-DNA haplogroups could not be observed thus adding Y-DNA haplogroups as covariates in a genetic association study is not essential (Figure 4). However there may be cases and cohorts where the contrary is true, thus an additional check on this can eliminate subtle population stratification due to non-recombining paternal ancestry of individuals within a sample.

**Figure 4:**
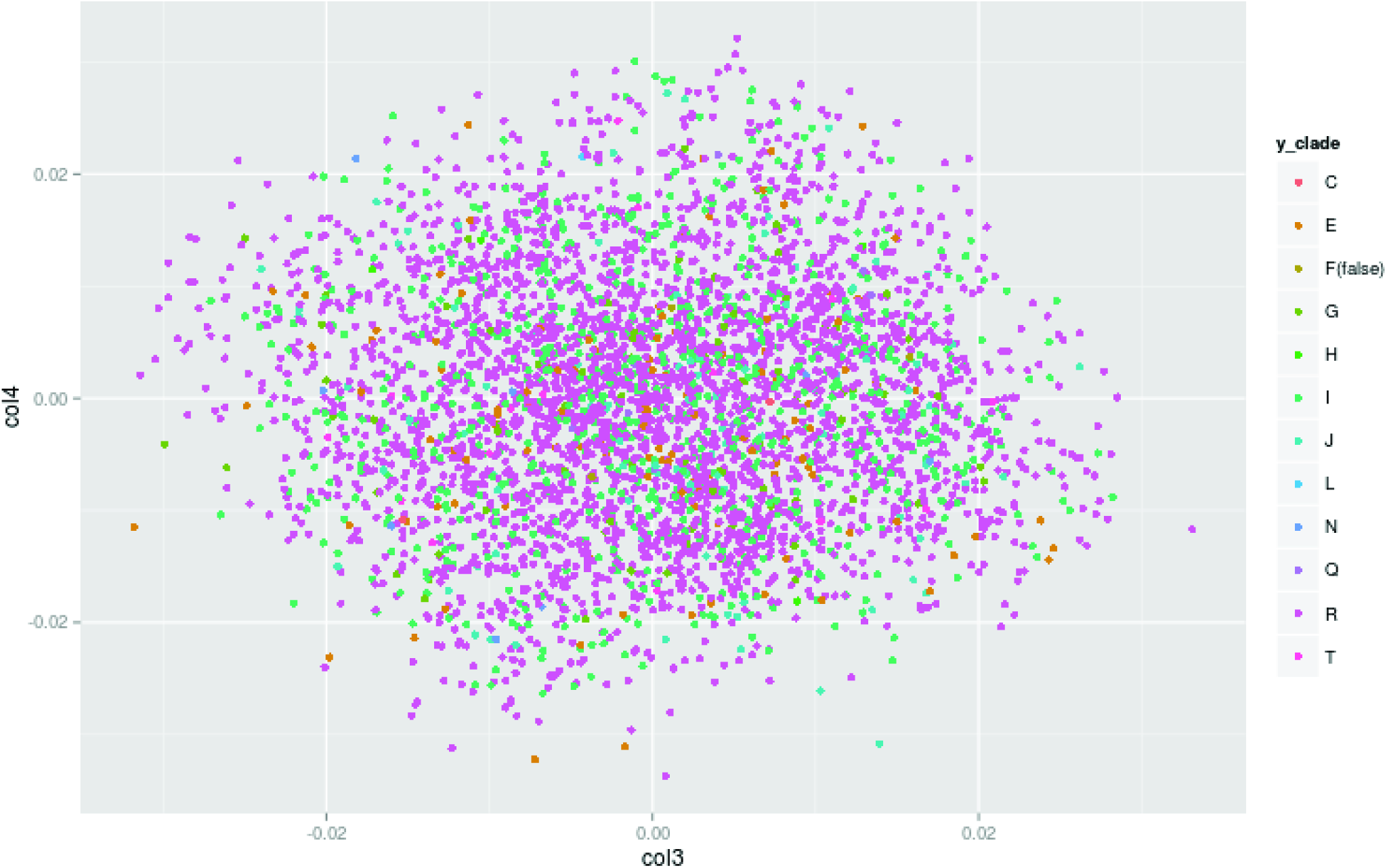
Y-DNA haplogroup vs top two principal components in ALSPAC individuals. Plotting the Y-DNA haplogroup clades on a PCA plot reveals that there is *no* apparent sub-clustering within the ALSPAC individuals. Thus adding Y-DNA haplogroup information as covariates to control for additional population stratification in ALSPAC is not needed. ggplot2 package in R was used to create the plot.

A substantial caveat of using Y-DNA haplogroups is the exclusion of females in the analyses. However, this limitation may be overcome with the combination of mitochondrial DNA haplogroup information. Another caveat is sample size; a problem for many European Y-DNA haplogroups, especially in the deeper sub-branches of the Y-DNA phylogenetic tree. The idea of using Y-DNA haplogroup information to inform genetic association studies is still underexplored and requires further research using different traits and haplogroups.

Overall, although structure could be a problem theoretically (and in some populations more than others), our results are in accordance with evidence showing that gross structure in common variant analysis does not seem to be a problem after PCA. On the other hand, recent evidence suggests that finer structure does exist for people of the British Isles [4]. It follows that if stratification is not really a problem, further studies could be efficiently improved by capturing some of this finer structure. This could be partially explained by variation of Y-DNA haplogroups.

Our result can be explained by chance and hence is not replicated. However, it illustrates the incorporation of Y-DNA haplogroups data which could be tested in Phenome-wide association studies (PheWAS) settings to systematically assess the impact of substructure.

## Acknowledgements

We are extremely grateful to all the parents and children who took part in this study, the midwives for their help in recruiting them and the whole ALSPAC team, which includes interviewers, computer and laboratory technicians, clerical workers, research scientists, volunteers, managers, receptionists and nurses. The UK Medical Research Council and the Wellcome Trust (Grant ref: 102215/2/13/2) and the University of Bristol provide core support for ALSPAC. ALSPAC is also part of the World Health Organisation initiated European longitudinal study of parents and children. This study makes use of data generated by the Wellcome Trust Case-Control Consortium and LabCorp (Laboratory Corportation of America) using support from 23andMe. A full list of the investigators who contributed to the generation of the data is available from www.wtccc.org.uk. Funding for the project was provided by the Wellcome Trust under award 076113 and 085475.

